# High-throughput AFM analysis reveals unwrapping pathways of H3 and CENP-A nucleosomes

**DOI:** 10.1101/2020.04.09.034090

**Authors:** Sebastian F. Konrad, Willem Vanderlinden, Wout Frederickx, Tine Brouns, Björn Menze, Steven De Feyter, Jan Lipfert

## Abstract

Nucleosomes, the fundamental units of chromatin, regulate readout and expression of eukaryotic genomes. Single-molecule experiments have revealed force-induced transient nucleosome accessibility, but a high-resolution unwrapping landscape in the absence of external forces is currently lacking. Here, we introduce a high-throughput pipeline for the analysis of nucleosome conformations based on atomic force microscopy and automated, multi-parameter image analysis. Our data set of ~10,000 nucleosomes reveals multiple unwrapping states corresponding to steps of 5 bp DNA. For canonical H3 nucleosomes, we observe that dissociation from one side impedes unwrapping from the other side, but unlike to force-induced unwrapping, we find only a weak sequence-dependent asymmetry. Centromeric CENP-A nucleosomes do not unwrap anti-cooperatively, in stark contrast to H3 nucleosomes, likely due to their shorter N-terminal α-helix. Finally, our results reconcile previously conflicting findings about the differences in height between H3 and CENP-A nucleosomes. We expect our approach to enable critical insights into epigenetic regulation of nucleosome structure and stability.

---

Nucleosomes are fundamental to the compaction of eukaryotic genomes into chromatin and function as regulators of gene activity^1–3^. While a large number of static nucleosome structures has become available in the past two decades^4,5^, the dynamic nature of nucleosomes^6^ and the role of epigenetic modifications^7^ remain unclear. Since dynamic structural changes influence the accessibility of nucleosomal DNA for readout^8^ and processing^9,10^, it is critical to understand nucleosomal unwrapping.

Canonical nucleosome core particles consist of two copies each of the four histones H2A, H2B, H3 and H4 assembled into a histone octamer that is tightly wrapped by ~147 bp of DNA^11,12^. The central ~121 bp of DNA contact structurally conserved histone-fold domains and the remaining ~13 bp of DNA at each end bind to the N-terminal alpha-helix^4^ (αN) of histone H3. Electrostatic interactions and hydrogen bonds stably pack the DNA onto the histone octamer, while DNA breathing, sliding, gaping, tightening, and loosening allow for nucleosomal dynamics on millisecond to minute time scales^13–16^. Partial unwrapping of the nucleosome core particle has been shown to occur anti-cooperatively^17^ with unwrapping on one end stabilizing the wrapped DNA on the opposite end in canonical nucleosomes^18^.

Numerous histone variants and post-translational modifications yield nucleosomal structures with varying degrees of stability and DNA wrapping^2,7,19^. In centromers –the chromosomal domains where both chromatids come together– H3 is replaced by the histone variant CENP-A, which has 64% sequence identity^20,21^ with H3. Crystallographic studies^22^ have revealed that in CENP-A nucleosomes the 13 bp of DNA at each end are more flexible than in H3 nucleosomes due to one missing helical turn of αN.

Atomic force microscopy (AFM) is a powerful tool to probe nucleosome structure and interactions due to its capability to image molecular complexes at the single molecule level label-free and with sub-nanometer resolution, well suited to visualize the DNA and protein components of nucleosomes^14,23–27^. Nevertheless, the accuracy and precision of measurements of structural parameters by AFM suffer from convolution of the molecular and AFM tip geometry, from the typically small sample sizes, and from inconsistencies associated with manual data analysis.

Here, we present an automated analysis pipeline for DNA and DNA-protein complexes in AFM topographic images that makes possible rapid and highly quantitative assessment of thousands of molecules with single-molecule resolution. Using the capabilities of our multi-parameter analysis, we reveal distinct unwrapping states of canonical H3 and CENP-A nucleosomes. We observe unwrapping of the two DNA ends to be anti-cooperative in H3 nucleosomes, consistent with previous reports. In contrast, no anti-cooperative unwrapping was found for CENP-A nucleosomes. Our results reconcile previously conflicting results on the height of CENP-A nucleosomes and reveal an important role of DNA crossovers in modulating the energy landscape of nucleosome wrapping.

## Results

### Automated AFM image analysis to quantify DNA and nucleosome conformations

We assembled nucleosomes by salt gradient dialysis on 486 bp DNA constructs under sub-stoichiometric conditions, such that the final sample contains bare DNA and predominantly mono-nucleosomes. Our DNA construct comprises a W601 nucleosome positioning sequence^28^ (147 bp) flanked by a short DNA arm (106 bp) and a long arm (233 bp) (Fig. 1a) and was used for both H3 and CENP-A nucleosomes (Methods). We deposited samples from aqueous buffer on a poly-L-lysine coated mica surface prior to rinsing and drying of the sample^29,30^. High-resolution images of the deposited nucleosome samples were obtained by amplitude modulation AFM in air (Fig. 1b). We developed an automated AFM image analysis pipeline to extract structural information of thousands of DNA and nucleosomes (Fig. 1c) and carried out multi-parameter analyses. Molecule detection consists of a background subtraction after applying a Gaussian filter and a subsequent skeletonization^31^ of both bare DNA (Fig. 1d) and nucleosomes (Fig. 1e). The skeletonized backbone of the molecules serves as the basis for molecule classification: whereas the skeleton of bare DNA contains exactly two endpoints and no branchpoints –points that have more than two neighbors– the skeleton of nucleosomes contains exactly two endpoints and two branchpoints. An adapted version of a previously published algorithm to trace DNA in AFM images^32^ measures the length of bare DNA molecules and nucleosome arms. Volume and height of the nucleosome core particle are estimated by fitting a half ellipsoid to the measured height data (Fig. 1e). The vectors connecting the DNA arm-ellipsoid intersections and the center of the ellipsoid define the nucleosome opening angle (Fig. 1e).

**Fig. 1.**
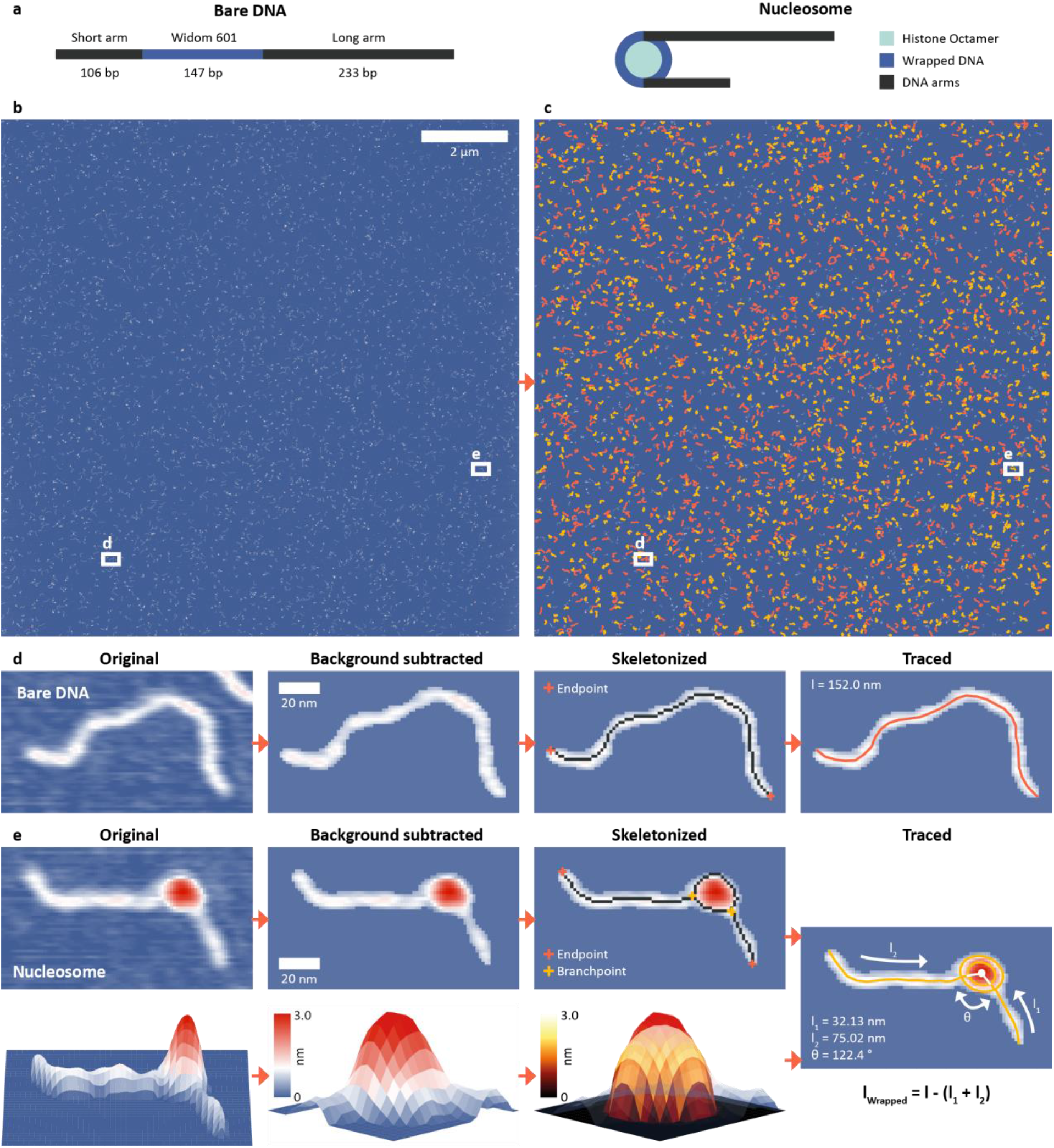
DNA and nucleosome structure parameters measured by automated AFM image analysis. **a**, Scheme of the DNA construct used throughout this work with a total length of 486 bp (106 bp + 147 bp Widom601 sequence + 233 bp) and the histone octamer (two copies each of H2A, H2B, H3 or CENP-A and H4). **b**, AFM image with a field of view of 12 μm x 12 μm with 8192^2^ pixels corresponding to a resolution of 1.46 nm/pixel. **c**, AFM image after tracing 1250 bare DNA strands (orange) and 1345 nucleosomes (yellow) with our automated image analysis pipeline. **d**, Image of a selected bare DNA strand: raw data, after background subtraction, after skeletonization, and after tracing. **e**, Image of a selected nucleosome: raw data, after background subtraction, and after skeletonization, together with a 3D surface plot of the nucleosome and the half ellipsoid fitted to the height profile of the core particle resulting in the traced nucleosome.

Our fully automated image analysis pipeline achieved a detection rate of ~95% (Supplementary Fig. 1), reducing the fraction of undetected molecules at least four-fold compared to the most advanced previous automated analysis framework for DNA-protein complexes^33^. Importantly, the automated analysis makes possible high-throughput analysis of AFM data obtained by high-speed large sample scanning: for example, imaging and automated tracing of a 12 *μm* x 12 *μm* field of view yielded structural parameters for 1250 bare DNA strands and 1345 nucleosomes (Methods and Fig. 1c).

### Identifying wrapping intermediates by multi-parameter analysis and image simulations

To quantify nucleosome wrapping, we first evaluated the average contour length of bare DNA molecules and found l_c_ = 152.9 ± 6.3 nm (mean ± std from 5651 molecules, Supplementary Fig. 1) corresponding to a length per bp of 0.314 ± 0.013 nm, in excellent agreement with previous measurements by AFM^34,35^ and solution X-ray scattering^36^. Similarly, we analyzed the DNA arm lengths of nucleosomes. By subtracting the combined nucleosome arm lengths from the contour length of bare DNA molecules, we obtain the distribution of wrapped lengths, i.e. the length of DNA confined in the nucleosome core particle. For a representative data set of H3 nucleosomes in buffer with 200 mM NaCl, we obtain a wrapped length of 135 ± 27 bp (mean ± std from 1011 molecules), in good agreement with previously reported values^33^. However, in contrast to previous reports^33,37,38^ we observed a bimodal distribution –rather than a single peak– for the wrapping of H3 nucleosomes (Fig. 2a). Fitting the wrapped length distribution to the sum of two Gaussians yields populations centered at 120 ± 14 bp and at 168 ± 12 bp. The distributions of opening angles (Fig. 2b) and of nucleosome core particle volumes (Fig. 2c) similarly suggest at least two different populations. However, the opening angle distribution is relatively flat, indicating that a large range of opening angles is sampled. To obtain a full quantitative view of nucleosome conformations, we exploit the fact that our analysis pipeline measures multiple parameters for each nucleosome particle to go beyond 1D distributions. Because of the solenoidal winding of nucleosomal DNA, we expect wrapped length and opening angle to be correlated and we indeed find that nucleosomes at wrapped lengths below 150 bp show a negative correlation between opening angle and wrapped length (Fig. 2d), suggesting that these nucleosomes populate states of partial unwrapping. The remaining nucleosomes at wrapped lengths between 160 bp and 190 bp (Fig. 2d) exceed the expected wrapping of the canonical nucleosome by ~20 bp.

**Fig. 2.**
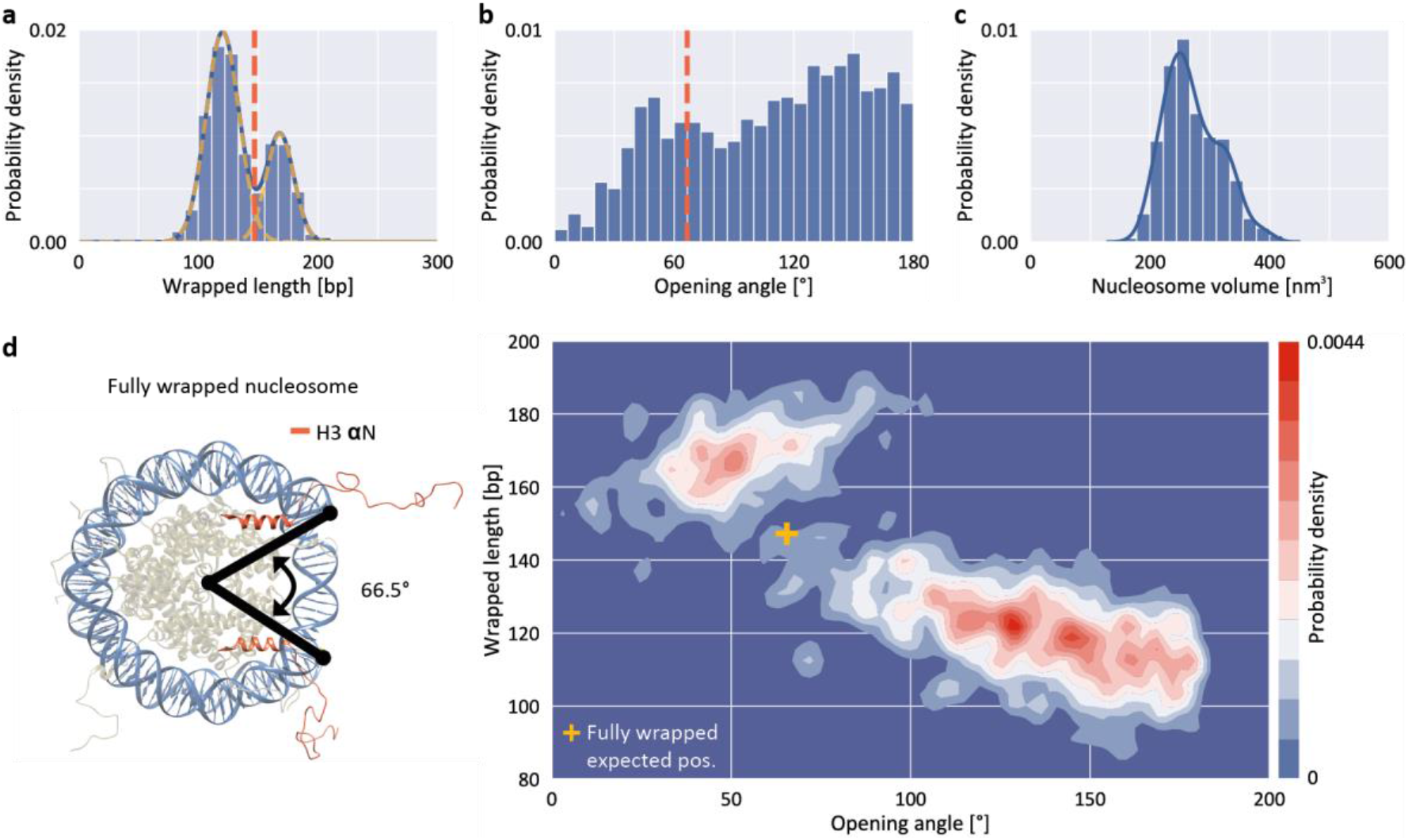
Structure parameters for H3 nucleosomes from AFM imaging. **a**, Wrapped DNA length distribution fitted using the sum of two Gaussians (centered at 120 ± 14 bp and 168 ± 12 bp). **b**, DNA opening angle distribution. The dashed orange line indicates the expected position of a fully wrapped nucleosome. **c**, Nucleosome core particle volume distribution. The solid line is a kernel density estimate. **d**, 2D kernel density profile (bandwidth = 2.5°, 2.5 bp) of nucleosome opening angle and wrapped length. The expected position of fully wrapped nucleosomes based on the crystal structure (left; rendered from PDB 1KX5) is shown as a yellow cross at an opening angle of 66.5° and a wrapped DNA length of 147 bp. The data set shown includes N = 1011 nucleosomes.

To quantitatively understand the observed 2D distributions, we simulated AFM images of nucleosomes with different levels of unwrapping. Simulated datasets were generated for eight states of unwrapping between 0 bp (fully wrapped; state 1) and 35 bp (state 8), with 5 bp wrapped length differences (Fig. 3a, Supplementary Fig. 2, and Methods), in line with the periodicity of the DNA helix, with results from single molecule DNA force spectroscopy experiments^39,40^, and with cryo-EM observations of nucleosome wrapping states^18^. After generating the ensemble of simulated nucleosome conformations, we applied AFM tip convolution and added experimental noise to create synthetic images that subsequently were analyzed using our automated framework (Fig. 3b). The slope of wrapped length vs. opening angle for measured nucleosomes at wrapped lengths below 150 bp (Fig 3c, −0.22 bp/°) agrees well with the slope predicted from simulated data (Fig. 3b, −0.23 bp/°), indicating that nucleosomes attach to the surface in a flat geometry with the DNA gyres parallel to the surface. However, we find that the analysis of the synthetic images systematically underestimates the opening angle by ~50° (mean squared error MSE = 20°) compared to the input configurations. This underestimation is the result of tip convolution in AFM imaging: due to the finite size of the AFM tip, the dimension of molecules is overestimated obscuring the exact entry/exit position of DNA in nucleosomes (Fig. 3d).

**Fig. 3.**
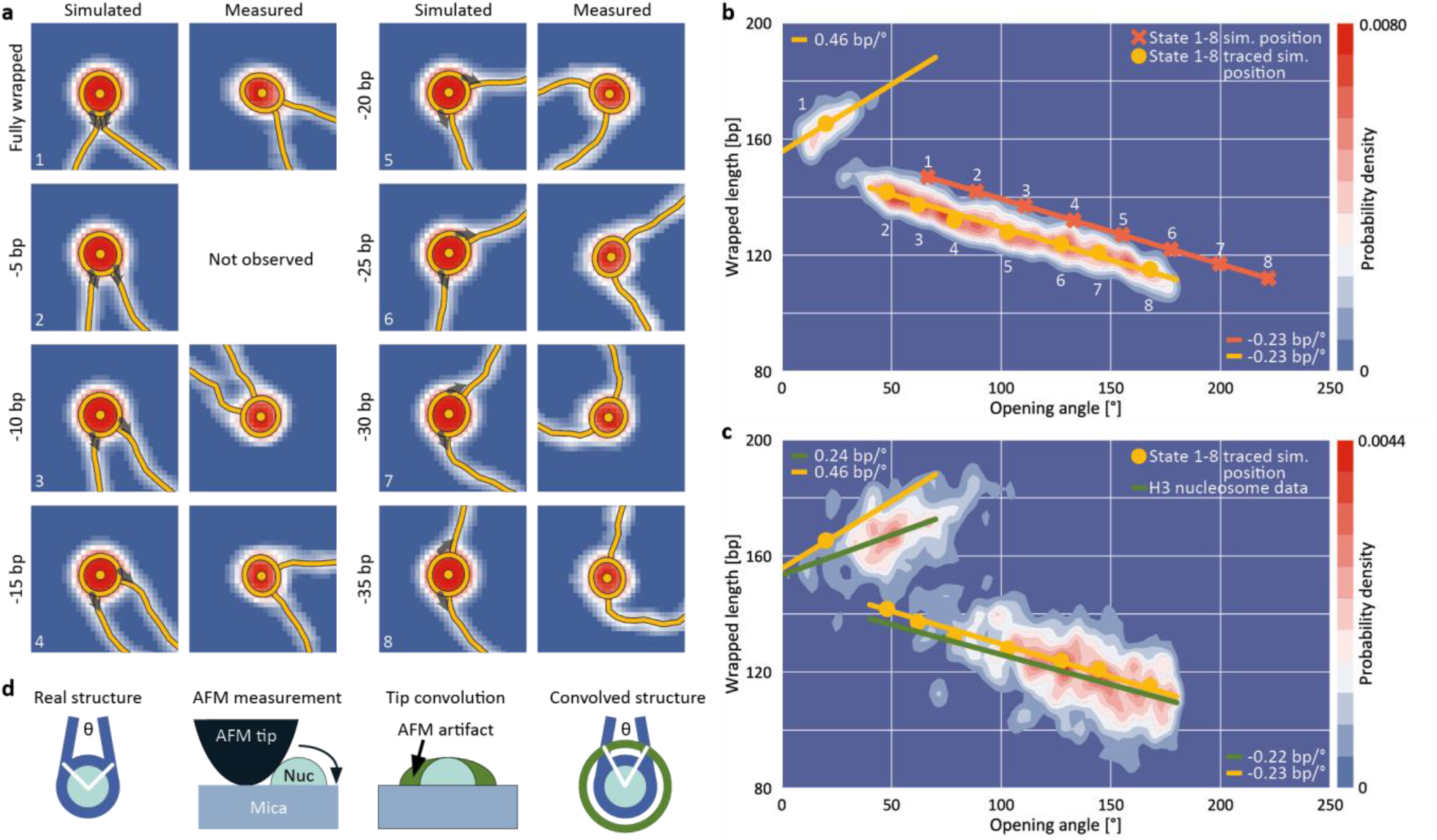
Assigning nucleosome wrapping states by comparison between simulation and experiment. **a**, Example synthetic nucleosome images of the eight simulated states and AFM imaged and traced nucleosomes with the same opening angle. The grey arrows indicate the starting direction of DNA simulation (Supplementary Fig. 2). For state 2, no measured counterpart was observed. **b**, 2D kernel density profile (bandwidth = 2.5°, 2.5 bp) of opening angle and wrapped DNA length for simulated nucleosome images (*N* = 1040, Supplementary Fig. 2). The simulations consist of eight states of nucleosome wrapping that differ by 5 bp of unwrapping. Orange crosses indicate the simulated positions based on the 5 bp unwrapping periodicity. Yellow circles indicate the centroids of each state found after analyzing the simulated images with our automated analysis pipeline. On average, the opening angle is shifted 50° (MSE = 20°) to lower angles due to the effect of AFM tip convolution. **c**, 2D kernel density profile (bandwidth = 2.5°, 2.5 bp) of opening angle and wrapped DNA length for AFM imaged nucleosomes (*N* = 1011). Regression lines fit to the experimental data (green) in comparison to the expected correlations (yellow) based on the simulations from (b). The top left population consists of fully wrapped nucleosomes in which the protruding DNA arms overlap in front of the nucleosome core particle whereas the lower population consists of nucleosomes in different states of DNA unwrapping. **d**, Sketch of AFM tip convolution resulting in an underestimation of the opening angle.

For the simulated partially unwrapped conformations (Fig. 3b, states 2-8; 5-35 unwrapped bp), the wrapped lengths determined from tracing of simulated images agree well with the input configurations (MSE = 4.2 bp). In contrast, the measured wrapped length for the simulated fully wrapped nucleosomes (state 1) exceeds the actual wrapped length of 147 bp by ~18 bp. This apparent wrapped length of 165 bp is fully consistent with the wrapped length of the second peak of the H3 nucleosome data (168 ± 12 bp; Fig. 2a,d, Fig. 3c). Our simulations rationalize why the apparent wrapped lengths for fully wrapped nucleosomes exceed the 147 bp expected from the crystal structure: the DNA arms that leave the nucleosome entry/exit site overlap close to the nucleosome core particle due to their initial directionality and the bending stiffness of DNA (Fig. 3a). AFM tip convolution obscures the crossing DNA strands and the software routine interprets the DNA crossover as being part of the nucleosome core, which in turn results in the apparent wrapped length values >160 bp. The simulated images of fully wrapped nucleosomes also reproduce the positive correlation between opening angle and wrapped length (Fig. 3b, state 1). Taken together, the results from simulated nucleosome structures strongly suggest that the population at ~165 bp wrapped length corresponds to fully wrapped nucleosomes (Fig. 3c, top left population), with the DNA arms crossing close to the nucleosome core particle.

### Opposing effects of salt concentration on nucleosome wrapping

While DNA is highly negatively charged, histone octamers carry significant net positive charge. Thus, histone-DNA interactions are sensitively modulated by ionic screening^41,42^. Additionally, crossing of the DNA arms exiting the nucleosome core particle presents an energy barrier that shapes the wrapping landscape. To investigate how ionic screening and DNA crossing at the nucleosome dyad affect nucleosome wrapping, we measured H3 nucleosomes deposited from buffer solutions of different ionic strengths ([NaCl] = 10 mM, 50 mM and 200 mM) and used our tracing software to generate 2D plots of wrapped length and opening angle (Fig. 4). To quantify the wrapping landscape of H3 nucleosomes as a function of ionic strengths, we first performed a principal component analysis (PCA) of nucleosome volumes and wrapped DNA lengths to separate fully and partially wrapped nucleosomes. The distributions of nucleosomes along the first principle component are bimodal for all data sets (Fig. 4a, insets). The first population corresponds to fully wrapped nucleosomes (white data points in Fig. 4a) and the second population to partially unwrapped nucleosomes (black data points in Fig. 4a). We find that the population of the fully wrapped state (31.2 %, 24.4 % and 13.8 % at 200 mM, 50 mM, and 10 mM NaCl, respectively; determined from the thresholds indicated in the insets of Fig. 4a) decreases with decreasing ionic strength, in line with increased like-charge repulsion of DNA at the exit/entry region at lower salt concentrations, as previously observed for divalent ions^38,43^ and for end-loops in supercoiled plasmids^44^.

**Fig. 4.**
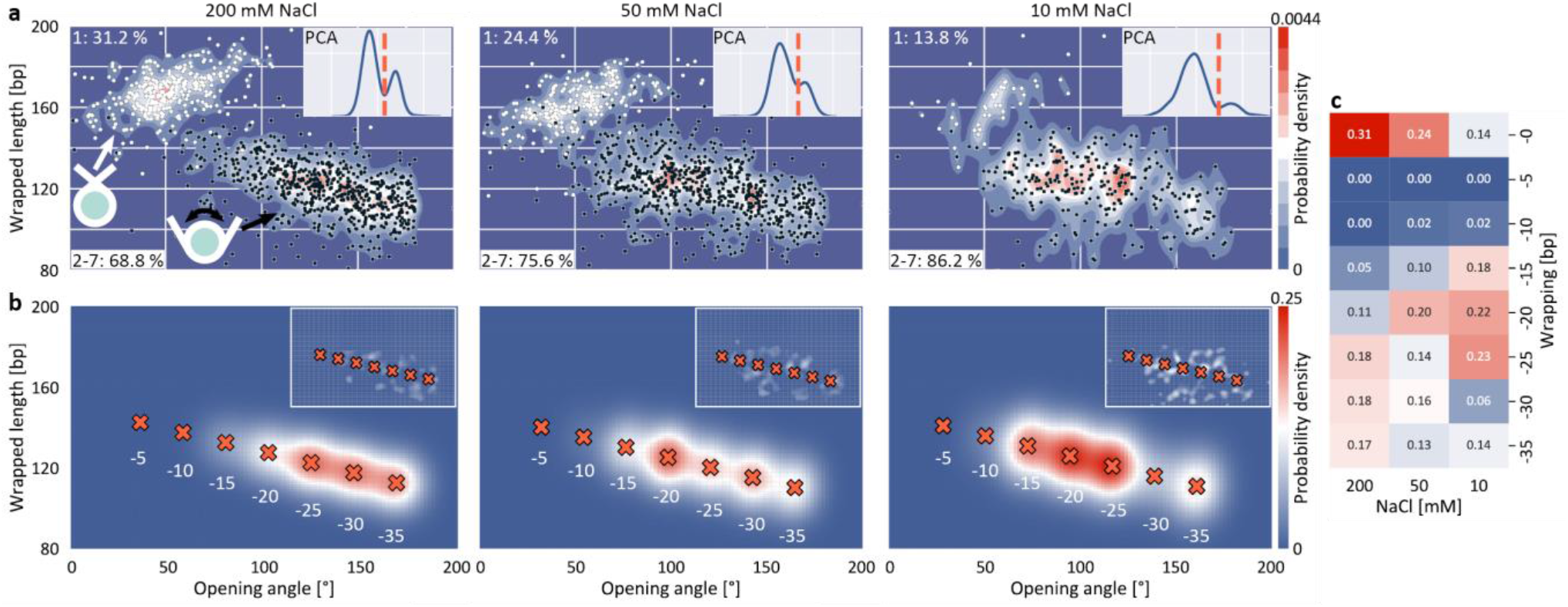
Effect of salt concentration on H3 nucleosome wrapping. **a**, 2D kernel density estimate profiles of wrapped length versus opening angle. Black and white dots indicate individual nucleosomes. The PCA of wrapped length and volume yields the basis for separating the two major populations as shown in the inset. The data sets include *N* = 1011 nucleosomes for 200 mM NaCl, *N* = 934 at 50 mM NaCl, and *N* = 325 at 10 mM NaCl. **b**, 2D Gaussian fits to the partially unwrapped nucleosomes. The Gaussian amplitudes yield the populations of the individual states of unwrapping and the insets show the fit residuals. 2D Gaussian fits to the partially unwrapped nucleosomes at 50 mM NaCl and 10 mM NaCl show the trend towards more compact wrapping for lower salt concentrations. All nucleosomes presented in this plot are from the same nucleosome reconstitution and were imaged with the same cantilever. **c**, Heat map of the populations of the individual wrapping states for NaCl concentrations of 200 mM, 50 mM and 10 mM.

To quantify how changing the ionic strength affects the distribution of the partially unwrapped states, we fitted seven 2D Gaussians located at fixed distances corresponding to the 5 bp periodicity and corrected for tip convolution based on the simulations. The amplitudes of the Gaussians represent the occupancy of the individual states of unwrapping and demonstrate a clear trend towards increased wrapping at reduced ionic strength, in line with increasing opposite-charge attraction that governs histone-DNA interactions and with previous work using FRET^45,46^. Together our data indicate two opposing trends on changing the ionic strength (Fig. 4c): decreasing salt concentrations decrease the occupancy of the fully wrapped state that features DNA crossing at the nucleosome dyad, but shift the distribution of partially unwrapped states towards less open conformations.

### Histone H3 nucleosomes unwrap anti-cooperatively

Building on the ability to precisely quantify nucleosome wrapping from high-throughput analysis of large AFM data sets, we next investigated the cooperativity in unwrapping behavior of H3 nucleosomes. The two sides of DNA exiting the nucleosome are distinguishable in our assay since the W601 sequence is placed asymmetrically, giving rise to a longer and a shorter DNA arm (Fig. 1a). The 2D distribution of short nucleosome arm length versus opening angle reveals a population at opening angles <80° and a bimodal distribution of short arm lengths for opening angles >80° (Fig. 5a). The population at opening angles <80° features short arm lengths of 75-95 bp, i.e. ~20 bp shorter as expected from the design of our DNA construct (106 bp), but consistent with the apparent length reduction due to the overlap of DNA at the dyad for fully wrapped nucleosomes, and can thus be assigned to the fully wrapped state. For opening angles >80°, i.e. the regime of partially unwrapped nucleosomes, the population splits into two branches, indicating that unwrapping can follow two distinct pathways. In the first pathway, the length of the short arm remains constant while the opening angle increases, suggesting exclusive unwrapping of the long arm. In the second pathway, the length of the short arm correlates positively with the opening angle (slope 0.20 bp/°) consistent with exclusive unwrapping of the short arm. The clearly separated pathways imply that unwrapping is anti-cooperative, i.e. that dissociation at one end suppresses unwrapping at the other. Our observation of anti-cooperative unwrapping is in agreement with previous reports based on single-molecule manipulation and FRET^17^ and on cryo-EM^18^, which revealed that unwrapping at one exit site stabilizes binding at the second exit site. Interestingly, a recent study modeling DNA caliper data found better agreement with a model where both arms can unwrap independently as compared to a model that includes anti-cooperativity^47^, in contrast to the clear anti-cooperativity visible in our and other previous data^17,18^.

**Fig. 5.**
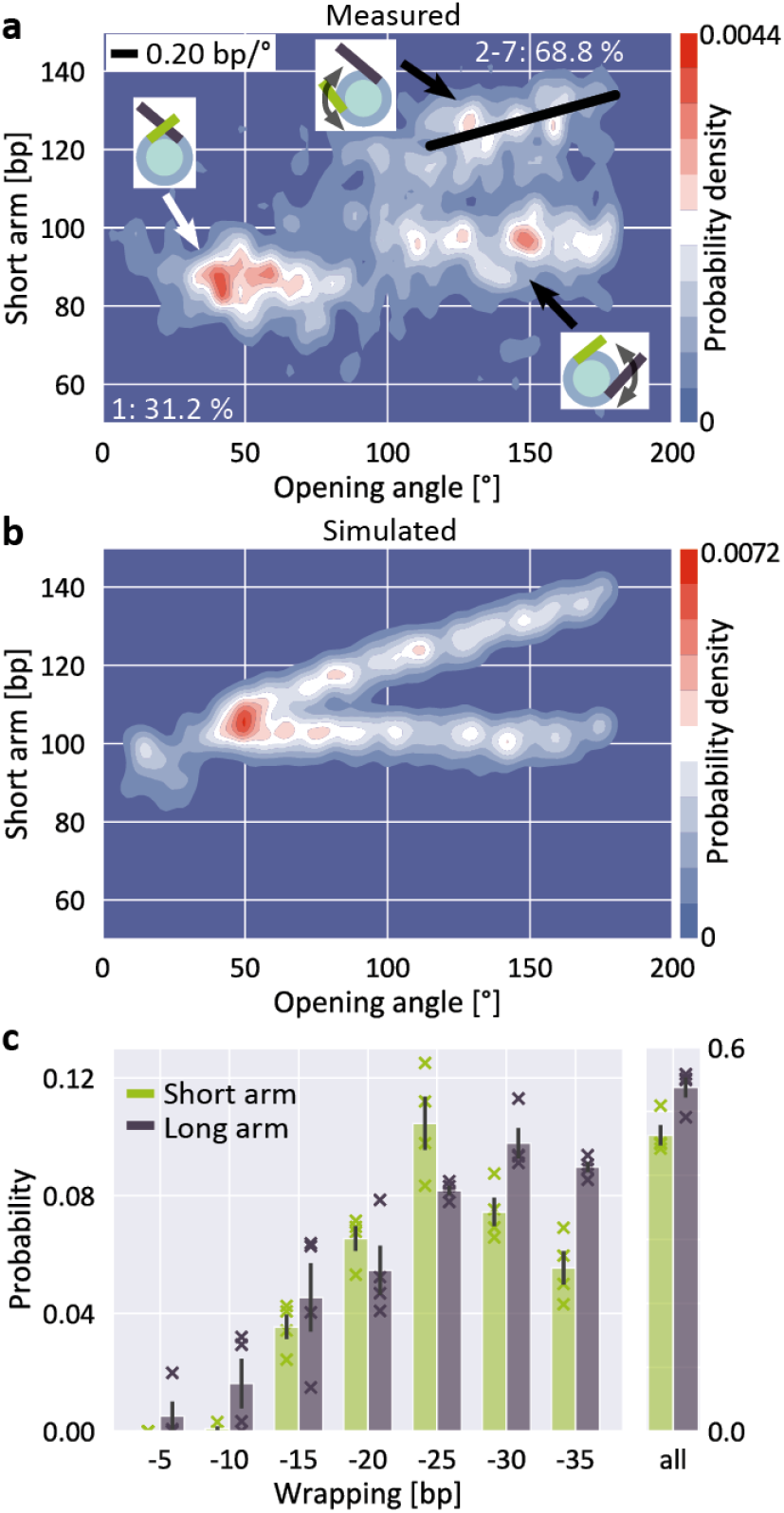
Anti-cooperative unwrapping of H3 nucleosomes. **a**, 2D kernel density profile (bandwidth = 2.5°, 2.5 bp) of short arm length and opening angle for H3 nucleosomes at 200 mM NaCl. A bimodal distribution for opening angles >80° is apparent, consistent with anti-cooperative unwrapping of the nucleosome core particle (*N* = 1011). **b**, 2D kernel density profile (bandwidth = 2.5°, 2.5 bp) of short arm length and opening angle for simulated nucleosomes (*N* = 1950). Unwrapping was simulated to occur either exclusively at the short arm or exclusively at the long arm, leading to a bimodal unwrapping behavior. **c**, 2D Gaussian fits to the density distribution of partially unwrapped nucleosomes (Supplementary Fig. 3) yield the populations of individual states of unwrapping. Unwrapping occurs significantly more likely (two-sample t-test *p* = 0.015) via the long arm (53.7 ± 1.6 %) than via the short arm (46.3 ± 1.6 %; mean ± SEM from four independent repeats) of the nucleosomes respectively. Data sets (*N* = 1011, 1524, 1480 and 815) comprise nucleosomes reconstituted in three independent nucleosome reconstitutions and imaged on two different AFM setups. Crosses indicate values from the individual data sets.

To quantify the propensity to unwrap via the distinct pathways, we simulated AFM images of nucleosomes featuring different levels of anti-cooperative unwrapping (Fig. 5b). Again, we fitted a linear combination of 2D Gaussians to the experimental density distribution of the partially unwrapped H3 nucleosomes using the expected positions based on the simulations to obtain population sizes along the different unwrapping pathways (Fig. 5c and Supplementary Fig. 3). We observed a small but significant (two-sample t-test *p* = 0.015) preference for long arm opening over short arm opening with probabilities of (53.7 ± 1.6) % and (46.3 ± 1.6) % respectively (mean ± SEM from four biological repeats). This preference for long arm opening reflects the non-palindromic nature of the W601 nucleosome positioning sequence: The DNA flanking the nucleosome dyad is less flexible on the long arm side compared to the short arm side (Supplementary Fig. 4) leading to the energetically more favorable unwrapping from the stiffer side, in line with previous force-induced experimental^17^ and computational^48^ unwrapping studies. However, in these studies nucleosomes almost exclusively unwrapped from the stiffer side, in contrast to the 54% to 46% partitioning that we observe. The difference might be caused by the experimental configurations: Force-induced nucleosome unwrapping with constrained DNA arms requires a nucleosome flip of 180° during unspooling and thus induces strong deformations in the DNA^48^. In contrast, thermal fluctuations and electrostatic interactions of the free DNA arms that drive nucleosome unwrapping in our study might be less susceptible to DNA flexibility. Quantifying the 5 bp unwrapping substates, we observe small, but significant differences in the unwrapping profiles for the two sides: unwrapping from the short arm side tends to occur by 20-25 bp, while the long arm side favors unwrapping by 30-35 bp (Fig 5c). These differences are in line with the free energy profile of W601 nucleosome unwrapping from a previous force-induced unzipping study^47,49^.

### CENP-A nucleosomes do not follow distinct unwrapping pathways

Previous Cryo-EM^20^ and AFM studies^50^ suggest that centromeric CENP-A nucleosomes exhibit enhanced structural dynamics and plasticity that deviates from canonical H3 nucleosomes. In contrast, magnetic tweezers measurements indicate that force-induced unwrapping and intrinsic stability of CENP-A and H3 nucleosomes are very similar^51^. To test CENP-A stability and unwrapping dynamics, we applied our AFM imaging and analysis pipeline to CENP-A nucleosomes.

The distribution of wrapped length and opening angle for CENP-A nucleosomes clearly differs from H3 nucleosomes under the same conditions (200 mM NaCl; compare Fig. 6a with Fig. 2d). First, only a small fraction of CENP-A nucleosomes populates the fully wrapped state with overlapping DNA arms (12.4 ± 1.7% for CENP-A vs. 27.0 ± 3.3% for H3, Fig. 6b,c). Second, CENP-A nucleosomes exhibit a shift of the partially unwrapped population towards more unwrapped states compared to H3 nucleosomes. Surprisingly, the clear negative correlation of wrapped length with opening angle for H3 nucleosomes is not apparent for CENP-A nucleosomes (Fig. 6a). We note that in our analysis, opening angles >180° are “folded back” and appear at smaller values (Fig. 6a, inset). To resolve higher unwrapping states, we examined again the first component of a PCA of volume and wrapped length and observed three populations (instead of two for H3 nucleosomes) separated by local minima (Fig. 6b, inset). One population corresponds to the fully wrapped state, as for H3. We assign the remaining two populations to partially unwrapped CENP-A nucleosomes with opening angles <180° and >180°, respectively. Correcting the opening angles of nucleosomes separated by the first local minimum of the first principal component led to the expected negative correlation of opening angle and wrapped length (Fig. 6b; −0.19 bp/°).

**Fig. 6.**
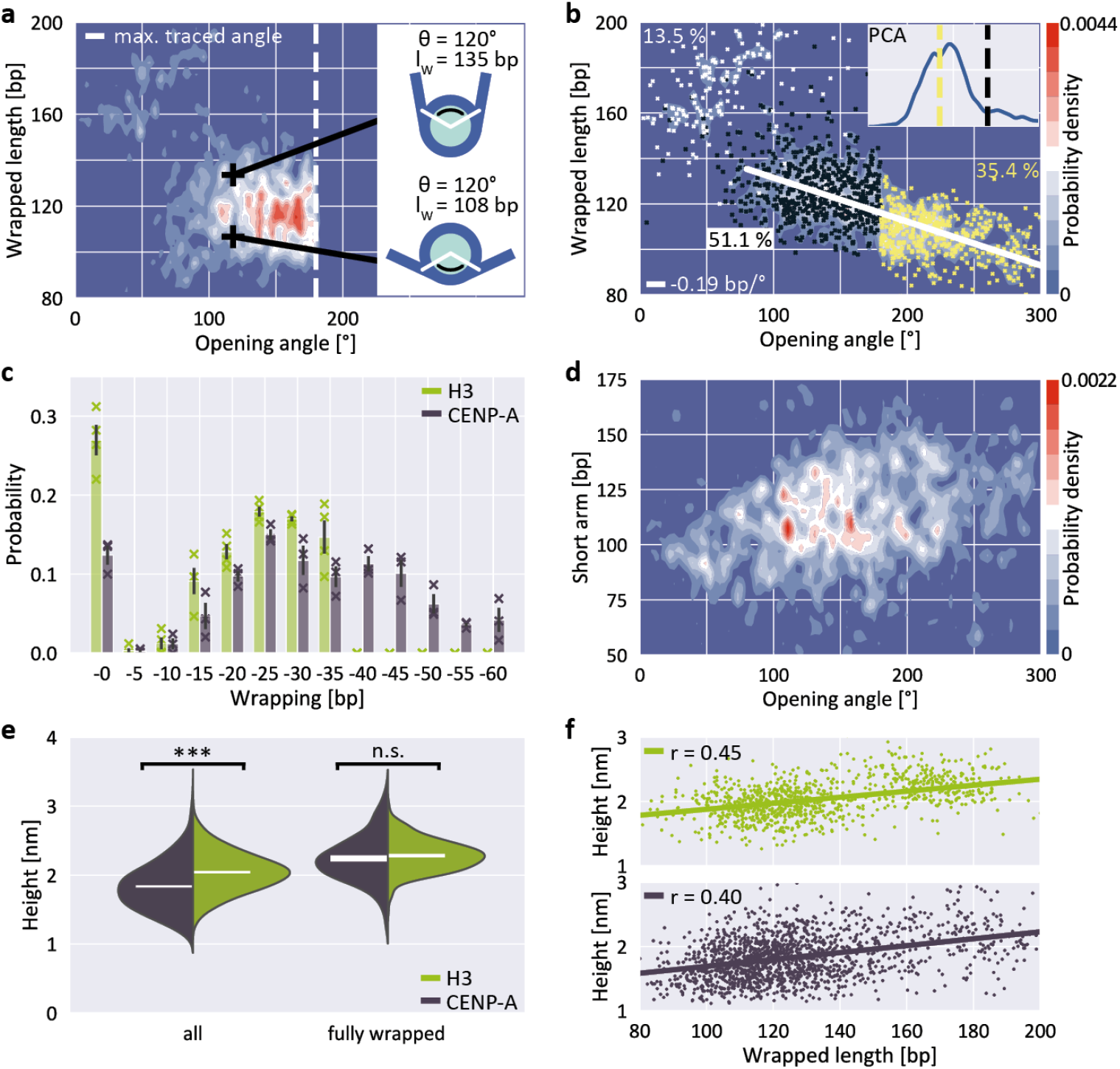
Unwrapping and heights of CENP-A nucleosomes. **a**, 2D kernel density profile (bandwidth = 2.5°, 2.5 bp) of wrapped length and opening angle for CENP-A nucleosomes (*N* = 1178, 200 mM NaCl). **b**, 2D kernel density profile (bandwidth = 2.5°, 2.5 bp) of wrapped length and opening angle for CENP-A nucleosomes after correcting the opening angles for folding back at 180°, based on the PCA of volume and wrapped length (inset). **c**, 2D Gaussian fits to the partially unwrapped CENP-A nucleosomes yield the populations of individual states of unwrapping (Supplementary Fig. 3). H3 populations are mean values obtained from four datasets (*N* = 1011, 1524, 1480 and 815) of nucleosomes from three biological repeats and imaged on two different AFM setups. CENP-A populations are mean values obtained from three datasets (*N* = 1178, 484 and 467) of nucleosomes from two biological repeats and imaged on two different AFM setups. **d**, 2D kernel density profile (bandwidth = 2.5°, 2.5 bp) of short arm length and opening angle for CENP-A nucleosomes indicating stochastic unwrapping of CENP-A nucleosomes in comparison to the anti-cooperative unwrapping of H3 nucleosomes. **e**, Violin plots of CENP-A and H3 nucleosome heights for the whole ensemble and for fully wrapped nucleosomes only. White bars are centered at the mean values of each distribution and thickness represents SEM. The height difference between CENP-A and H3 is highly significant for all nucleosomes (two-sample t-test *p* = 1.6·10^−60^) and not significant for fully wrapped nucleosomes (*p* = 0.11). **f,** Correlation between wrapped length and nucleosome height for H3 (top) and CENP-A nucleosomes (bottom).

Fitting the populations of unwrapping states by 5 bp steps in the CENP-A nucleosome data (Fig. 6c), we find that the most frequented states of CENP-A nucleosomes lie between 25 bp and 35 bp unwrapping, similar to H3 nucleosomes under the same conditions (Fig. 4c). However, in contrast to H3 nucleosomes, CENP-A nucleosomes also significantly populate unwrapping states between 40 bp to 60 bp (35.4 ± 4.6% of the total population).

Third, in addition to the overall shift to less wrapped states, our CENP-A nucleosome data reveal a striking difference to H3 nucleosomes in wrapping pathways: While the H3 nucleosome data feature a clearly bimodal distribution of short arm length vs. opening angle for angles >80° (Fig. 5a), indicative of anti-cooperative unwrapping, the CENP-A data exhibit no such branching and a broad distribution of short arm lengths instead (Fig. 6d), suggesting that unwrapping of the two arms is not anti-cooperative for CENP-A nucleosomes.

Recent Cryo-EM work^18^ has suggested that DNA unwrapping on one side triggers a conformational change of the adjacent H3 αN, which in turn leads to rearrangement of H3 αN on the opposite side, resulting in stabilization of the DNA contact on the wrapped side. Together, these allosteric changes constitute a latch mechanism that likely contributes to the asymmetric unwrapping of H3 nucleosomes. In CENP-A nucleosomes the αN helix is shortened^20^ compared to H3. The lack of anti-cooperative unwrapping revealed by our data suggest that the reduced αN helix in CENP-A nucleosomes is insufficient for the latch mechanism, leading to stochastic unwrapping of DNA from CENP-A nucleosomes from both sides, in line with a recent molecular dynamics study^52^. Interestingly, we observe that the most unwrapped configurations (Fig. 6d, opening angles >200°, corresponding to >40 bp unwrapped) consistently involve unwrapping of both DNA arms, giving rise to intermediate short arm lengths. This is consistent with the observation that unwrapping of >40 bp, i.e. opening angles >200°, from one side leads to disruption of the overall nucleosome structure^18^ and, therefore, nucleosomes with unwrapping of >40 bp exhibit concurrent unwrapping on both sides.

### Differences in DNA wrapping reconcile previously conflicting results on CENP-A nucleosomes

A previous AFM study proposed that CENP-A nucleosomes have a reduced height compared to H3 nucleosomes^53^. However, this finding was subsequently challenged and follow up studies found no height difference for CENP-A and H3 nucleosomes^54–56^. In our measurements we find that overall H3 nucleosomes are significantly higher than CENP-A nucleosomes (two-sample t-test *p* = 1.6·10^−60^; Fig. 6e), with mean heights of (2.04 ± 0.01) nm for H3 (mean ± SEM from *N* = 1011 molecules) and (1.83 ± 0.01) nm for CENP-A nucleosomes (*N* = 1645 molecules). Our results are in agreement both regarding the significant height difference between H3 and CENP-A nucleosomes and in terms of the absolute height values with the first study that reported the height differences^53^. However, we observe a significant correlation between wrapped length and measured height (Fig. 6f; Pearson’s *R* = 0.45 and *p* = 7.1 · 10^−52^ for H3 nucleosomes and *R* = 0.40 and *p* = 2.4 · 10^−65^ for CENP-A nucleosomes). Since CENP-A nucleosomes are on average less wrapped (Fig. 6c), the correlation between wrapping and height implies that differences in wrapping might account for the reduced height of CENP-A nucleosomes. Comparing only fully wrapped H3 and CENP-A nucleosomes (classified based on the PCA results, see Fig. 4a and Fig. 6b), we indeed find mean heights of (2.28 ± 0.01) nm (*N* = 315 molecules) and (2.24 ± 0.02) nm (*N* = 205 molecules), respectively, corresponding to no significant difference in the mean heights (two-sample t-test *p* = 0.11).

In summary, we find that if we average over all wrapping states, CENP-A nucleosomes are lower than H3 nucleosomes, in agreement with an initial report of nucleosome height difference in AFM^53^. However, if we only consider fully wrapped nucleosomes, we do not detect a significant difference, similar to follow up work motivated by the initial study^54,55^. The apparently conflicting findings might be explained by differences in imaging conditions. In particular, the three studies used different salt conditions (10 mM NaCl and 1 mM EDTA vs. 150 mM NaCl + 2 mM MgCl_2_ vs. 1 mM EDTA only)^53–55^, which can significantly alter unwrapping (Fig. 4). In addition, they used different underlying DNA constructs (nucleosome arrays on linear DNA vs. nucleosome arrays on supercoiled plasmids vs. mononucleosomes on short DNA), which also might affect wrapping. We conclude that in addition to precisely controlling imaging parameters, it is important to compare AFM measured heights for nucleosomes occupying the same state of wrapping, highlighting the power of our multi-parametric analysis approach.

## Discussion

Quantitative assessment of nucleosome conformations is a key to understanding regulation of DNA accessibility and the binding of transcription factors^8^. In this work, we introduce an automated framework that enables high throughput analysis of AFM images of nucleoprotein complexes and applied it to canonical H3 and centromeric CENP-A nucleosomes. By exploiting correlations between different structural parameters of ~10,000 nucleosomes, we map molecular ensembles along different degrees of freedom, which in turn allows us to extract detailed nucleosome wrapping landscapes. We use simulations of AFM images to understand how tip-convolution in AFM imaging affects the observed structure parameters and to quantify the occupancy of different states of wrapping from our experiments. We observed partial unwrapping of ~70% of the canonical H3 nucleosomes at close to physiological salt concentrations ([NaCl] = 200 mM) in agreement with previous electron microscopy^43^, AFM^38^, and solution SAXS^57^ studies of nucleosomes. In fully wrapped nucleosomes, the DNA arms overlap in close proximity to the DNA entry/exit region of the nucleosome core particle. DNA crossover at the exit region presents a significant energy barrier that might regulate nucleosomal DNA readout either by binding of histone H1 to form repressive chromatosomes or by granting access to RNA polymerases^8^ and molecular machines that process the genetic code.

Our data demonstrate pronounced anti-cooperative unwrapping of H3 nucleosomes and preferential unwrapping from the stiffer side of the non-palindromic W601 sequence in agreement with previous single-molecule and cryo-EM studies^17,18,49^. However, our data show only a slight preference for unwrapping from the stiffer side in contrast to previous studies that have seen it almost exclusively. While our methodology captures a molecular ensemble in thermal equilibrium, previous results were obtained via force-induced unwrapping or for constructs containing only 147 bp of DNA^18,57^, which might account for the differences. Both experimental approaches are of physiological relevance since nucleosomes can be invaded either passively due to spontaneous fluctuations^58^ or actively by forces generated by polymerases and chromatin remodelers^59,60^. We speculate that our approach samples the clearly distinct nucleosomal unwrapping landscape for passive invasion of nucleosomes with linker DNA in contrast to previous force-induced unwrapping assays.

In contrast to their canonical H3 counterparts, we find centromeric CENP-A nucleosomes to be substantially less wrapped than H3 nucleosomes, with 2-fold fewer nucleosomes being fully wrapped. This result is in agreement with previous high-resolution structural studies by X-ray crystallography^22^ and cryo-EM^20^ that assigned the shortened N-terminal α-helix of histone H3 to weakened interaction with DNA at the exit points of the nucleosome. More than 30% of the population of CENP-A nucleosomes unwrapped >35 bp. Unwrapping >40 bp requires partial opening of both arms, which is suppressed in H3 nucleosomes. Our data are in line with results from force-induced unzipping of nucleosomal DNA: peak-forces associated with the strong barrier between superhelical locations 3.5 and 6.5, i.e. up to 40 bp unwrapping, were significantly reduced in yeast centromeric versus H3 nucleosomes^39^. However, DNA-histone interactions closer to the dyad were equally strong for yeast centromeric and H3 nucleosomes preventing unwrapping of more than 40 bp from one side.

Our data show no anti-cooperativity in the unwrapping of CENP-A nucleosomes, in contrast to H3 nucleosomes. We propose that the stabilizing latch mechanism that contributes to anti-cooperativity^18^ in H3 nucleosome unwrapping is missing in CENP-A nucleosomes due to the shortened αN helix. Since CENP-A is a key epigenetic mark to maintain structural integrity of the centromer, we speculate that both stochastic unwrapping and overall decreased DNA wrapping of CENP-A nucleosomes might facilitate binding of proteins to specific DNA sequences in the centromer. For example, the centromer-DNA binding protein complex CBF3 is essential for chromosome segregation and binds selectively to the highly conserved CDEIII DNA sequence found in centromers^61^.

In summary, we have developed a high-throughput automated analysis platform and used it to uncover thermally activated pathways of H3 and CENP-A nucleosome wrapping in unprecedented detail. Our methodology will facilitate future (high-speed) AFM studies that involve structure and interactions of nucleoprotein complexes by either using fast imaging of large molecular ensembles or by time-lapsed imaging of the molecular dynamics at the single molecule level^27^.

## Acknowledgements

We thank Philipp Korber and Felix Müller-Planitz for help with initial nucleosome reconstitutions, Herlinde De Keersmaecker for assistance with AFM imaging, and Thomas Nicolaus for help with sample preparation. This work was funded by the Deutsche Forschungsgemeinschaft (DFG, German Research Foundation) through SFB863 – Project ID 111166240. W. F., T. B., and S. D. F. acknowledge KU Leuven – internal funds (IDO) and Fund for Scientific Research (FWO).

## Author contributions

S.F.K., W.V. and J.L. designed the experiments. S.F.K. performed biochemical experiments and W.F., T.B. and S.D.F. supported AFM measurements. B.M. assisted with data analysis. S.F.K., W.V. and J.L. wrote the paper with input from all authors.

## Competing interests

The authors declare no competing interests.

## Methods

### DNA purification and nucleosome reconstitution

DNA was PCR amplified from a GeneArt High-Q String DNA fragment (Thermo Fisher) containing the Widom 601 positioning sequence. The DNA was purified using a QIAquick PCR purification kit (Qiagen) and subsequently eluted to a volume of 30 μL with ddH_2_O. Histone proteins were purchased from EpiCypher. While the H3 histones were available as part of recombinant human histone octamers, CENP-A histones were purchased as CENP-A/H4 tetramers and added to the dialysis chamber together with an equimolar ratio of H2A/H2B tetramers. Nucleosome reconstitution was performed via salt gradient dialysis^62^. The dialysis chamber contained 0.65 μg of the histone octamers and 3 μg of the 486 bp DNA at 2 M NaCl and was placed in one liter of high-salt buffer at 2 M NaCl. Over a course of 15 hours, three liters of low-salt buffer at 50 mM NaCl were transferred to the high-salt buffer at 4°C. Finally, the dialysis chambers were moved to one liter of low-salt buffer for three hours.

### AFM sample preparation and imaging

The sample mix containing bare DNA and reconstituted nucleosomes –usually 30% to 50% of the DNA strands do not bind to histones based on the individual concentrations used during the nucleosome reconstitution– was incubated at the desired salt concentration (10 mM NaCl/50 mM NaCl/200 mM NaCl and 10 mM Tris-HCL, pH 7.6, for all measurements) for 1 min on ice. The solution was then deposited on a freshly cleaved poly-L-lysine (0.01% w/v) coated muscovite mica for 30 seconds and subsequently rinsed with 20 mL of milliQ water before drying with a gentle stream of filtered N_2_ gas.

We used two different commercial AFM instruments for imaging. All AFM images were acquired in tapping mode at room temperature. One set of images was acquired on a Multimode VIII AFM (Bruker) using silicon tips (AC160TS, drive frequency of 300-350 kHz, Olympus). Images were scanned over a field of view of 3 μm x 3 μm at 2048 x 2048 pixels with a scanning speed of 1 Hz. Independent measurement repeats were performed on a Nanowizard Ultraspeed 2 (JPK) with reflex gold coated tips (USC-F5-k30, drive frequency 5000 kHz, Nanoworld). Here, images were scanned over a field of view of 6 μm x 6 μm at 4096 x 4096 pixels with a scanning speed of 3 Hz or over a field of view of 12 μm x 12 μm at 8192 x 8192 pixels at 3 Hz (Fig. 1b). For H3 nucleosomes, four data sets were acquired at 200 mM NaCl and 10 mM Tris over three separate nucleosome reconstitutions. For CENP-A nucleosomes, three data sets were acquired at 200 mM NaCl and 10 mM Tris over two separate nucleosome reconstitutions.

### AFM image analysis

We developed an automated image analysis pipeline to analyze the flattened AFM images. For the AFM data analyzed in this work, a background height threshold of 0.16 nm was applied. The bare DNA strands were traced with 5 nm segments^32^ from both sides separately and the mean value was used as contour length. Over 95% of the viable molecules in the images were detected automatically. Here, viable molecules are defined as molecules that do not have overlaps with other molecules and can be analyzed by manual tracing. To achieve an even higher detection rate, manual input allowed the separation of unclassified objects (for example for two DNA arms that slightly overlap and thus prevent automated detection). This way, 98% of all viable nucleosomes of the example image were detected (Supplementary Fig. 1). Even with manual help for detecting and classifying individual molecules, all measured and presented structure parameters were obtained by the structure analysis routine of the toolbox. The four H3 data sets at 200 mM NaCl consist of 1011, 1524, 1480 and 815 analyzed nucleosomes. The three CENP-A datasets consist of 1178, 484 and 467 analyzed nucleosomes.

### AFM image simulations

Fully wrapped nucleosome images were simulated by creating a disk with a diameter of 11 nm and uniform height and simulating 2D worm-like chains with lengths of 233 bp and 106 bp that protrude the disk at an opening angle of 66.5°. The direction of the DNA chains was deduced from the crystal structure of the canonical nucleosome (PDB 1KX5, Fig. 2d, Supplementary Fig. 2). Consecutively, the DNA was dilated to its expected width of 2 nm and random noise in combination with a Gaussian filter (*σ* = 2 nm) was applied to mimic the effect of tip convolution. Partially unwrapped nucleosomes were simulated by adding base pairs to one end of the simulated chains in 5 bp steps, increasing the opening angle by 4.45° per base pair of unwrapping (based on 147 bp wrapped over a total of 654° in the crystal structure) and adjusting the direction of the protruding DNA arms. Similarly, synthetic images of bare DNA were simulated with a 2D worm-like chain of 486 bp and the same steps of dilation and tip convolution. The synthetic bare DNA images were analyzed for their average DNA contour length that is needed to calculate the wrapped length in nucleosomes in our automated readout pipeline. The simulated images were analyzed with the same automated readout software as the experimental images.

## Data availability

Data supporting the findings of this manuscript are available from the corresponding authors upon reasonable request.

## Code availability

Code is available on https://github.com/SKonrad-Science/AFM_nucleoprotein_readout including a detailed installation guide and an example image.

## Notes

### Competing Interest Statement

The authors have declared no competing interest.

