## Supplementary Information for "High-throughput AFM analysis reveals unwrapping pathways of H3 and CENP-A nucleosomes"

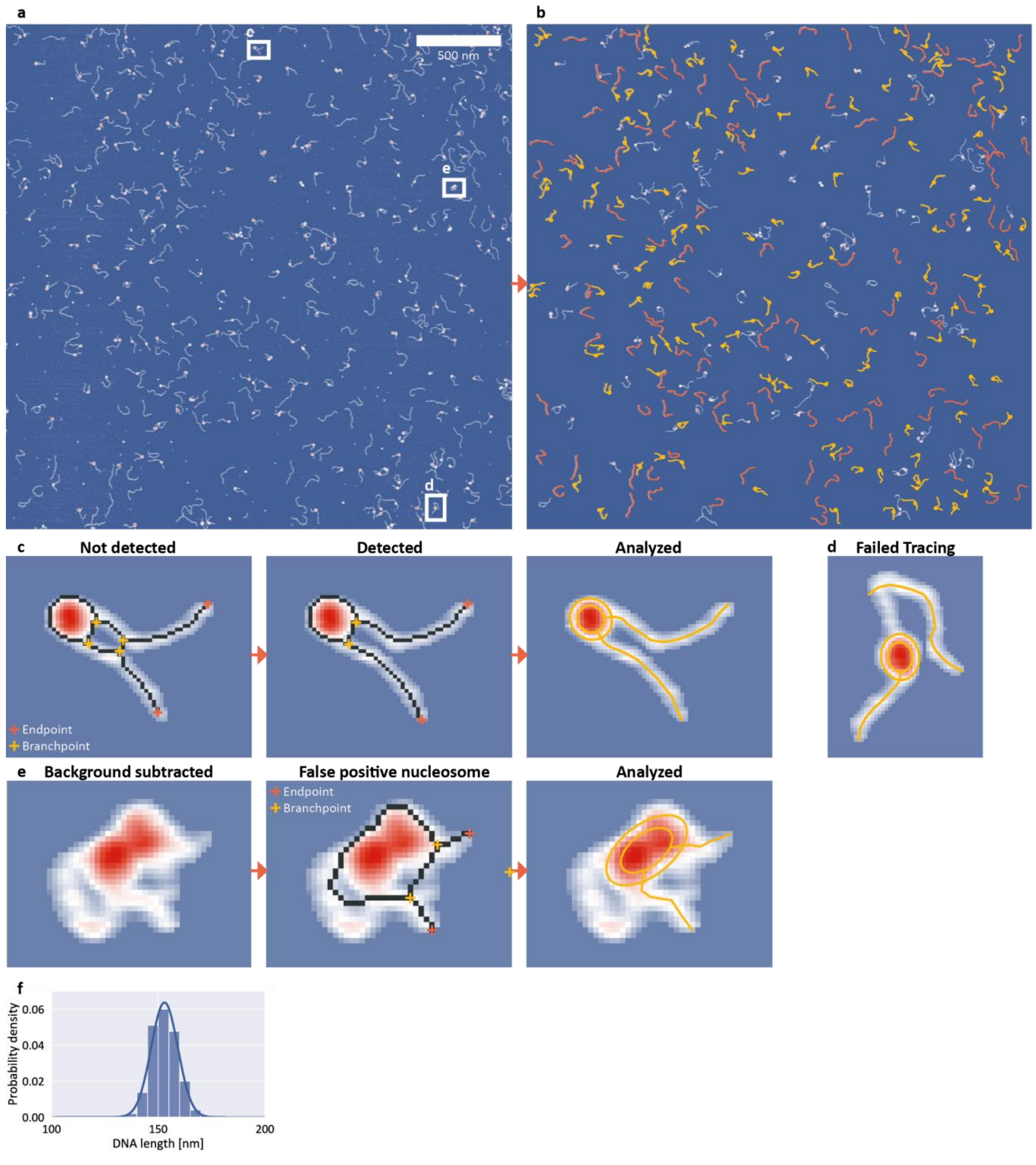


**Supplementary Fig. 1 | Detection efficiency, manual molecule classification, and DNA length**.

**a**, AFM topographic image of bare DNA and nucleosomes to assess the detection rate of the automated tracing. The total field of view is 3 µm x 3 µm and was recorded with 1.46 nm/pixel.

**b**, Analysis of the topographic image from panel **a**. 112 bare DNA strands (orange) and 131 nucleosomes (yellow) were detected and analyzed automatically with manual detection help as described in **c**.

To quantify the detection rate and to assess imaged molecules that were not classified as either DNA or nucleosome in the first fully automated step, the image was inspected visually. The fully automated classification and tracing routine detected 95% of all manually analyzable DNA and nucleosomes in the field of view that did not show overlaps with other molecules.

**c**, Example how manual intervention can improve classification of molecules not assigned to bare DNA or nucleosome molecules in the first automatic classification step. In the case depicted here, too many branchpoints (four instead of two) are detected and thus the nucleosome was not automatically classified. After manually removing the overlapping pixels of the DNA arms, the nucleosome is properly classified and the automated analysis framework traces the structure parameters. Overall, such manual classification help for unclassified molecules enabled tracing of up to 98% of manually analyzable molecules.

**d**, Example of a false negative, i.e. a molecule that by visual inspection appears to constitute a valid nucleosome or DNA, but is not traced properly. The tracing fails due to the strong bending of the long arm. Overall, 2% of molecules that we identified as valid by visual inspection fall into this category.

**e**, Example of a false positive, i.e. a molecule that is classified as a valid nucleosome or DNA by our algorithm, but excluded by visual inspection. In the image shown in **a**, a total of 9 false positive molecules were traced by the automated toolbox. For this work, false positives were removed for further analysis. The example molecule contains two branchpoints and two endpoints and is thus classified as a nucleosome.

**f**, Histogram of bare DNA lengths of all data sets measured in 200 mM NaCl presented in this work. We find a contour length of $l_c = 152.9 \pm 6.3$ nm (mean ± std from 5651 molecules).

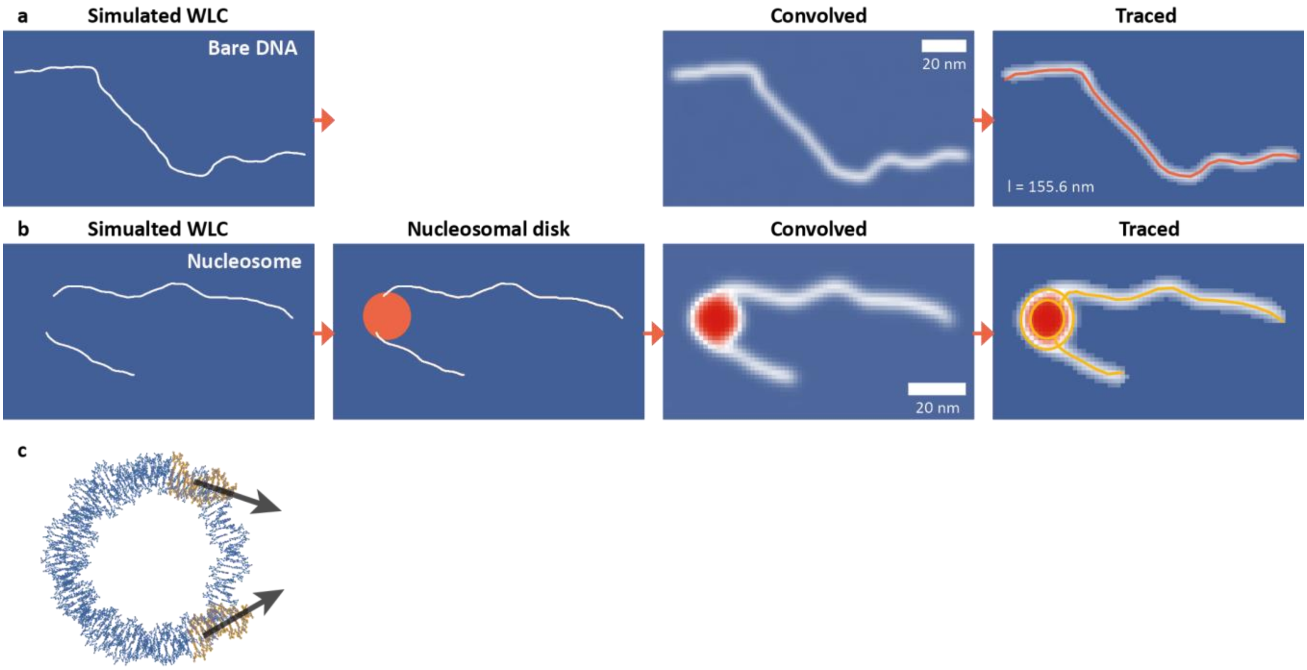


**Supplementary Fig 2 | DNA and nucleosome simulations to quantify the effect of AFM tip convolution.**

**a**, 486 bp DNA were simulated using the worm-like chain polymer model with a persistence length of 40 nm and a rise per base pair of 0.32 nm/bp. The simulated DNA backbone was dilated to a width of 2 nm and convolved with a Gaussian filter after applying random noise to mimic the effect of AFM tip convolution. The simulated DNA strands were then traced with our automated analysis pipeline using the same settings as used for experimental data. See Methods for details.

**b**, Simulation of nucleosomes consisted of generating two DNA arms (106 bp and 233 bp for the fully wrapped nucleosome) based on the worm-like chain model that protrude from the nucleosomal disc. The orientation of the protruding DNA arms was deduced from the crystal structure (panel **c**), for details see the section "AFM image simulations" in Methods. The nucleosome depicted here is unwrapped by 35 bp from the long arm side.

**c**, The crystal structure of the nucleosome core particle yields the orientation of the DNA arms and the nucleosomal opening angle of 66.5° for fully wrapped nucleosomes. Rendered from PDB 1KX5. For partially unwrapped nucleosomes the DNA orientation and opening angle are adjusted accordingly (Methods).

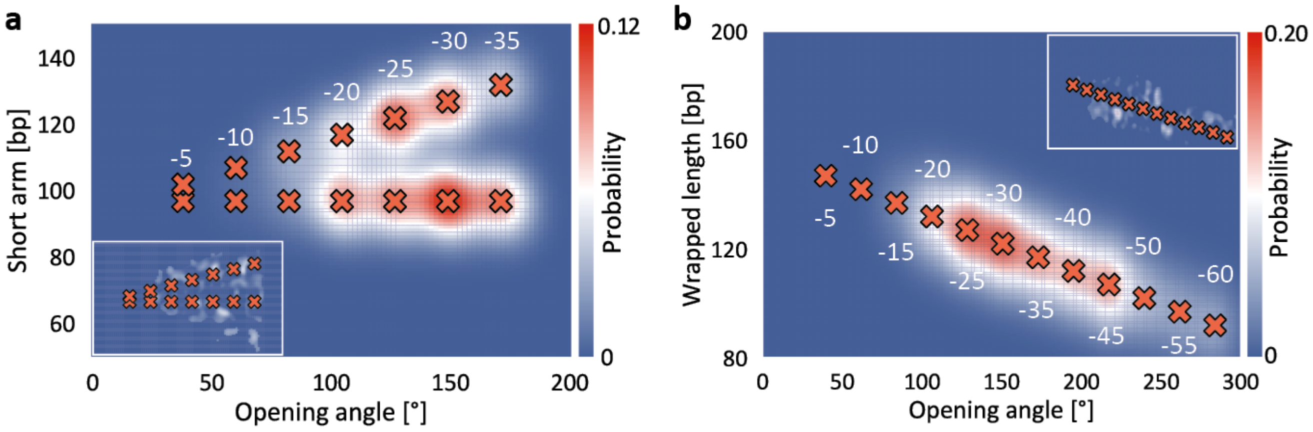


**Supplementary Fig 3 | Determination of the occupancies of different wrapping states.**

**a**, Fit of 14 2D Gaussians to the density distribution of short arm length and opening angle for the partially unwrapped H3 nucleosomes (Fig. 5a) measured at 200 mM NaCl (N = 696). The inset represents the difference between the measured and the fitted 2D density profile, i.e. the residuals of the fit. The occupancies determined from the fits are shown in Fig. 5c.

**b**, Fit of 12 2D Gaussians to the density distribution of wrapped length and opening angle for the partially unwrapped CENP-A nucleosomes (Fig. 6b) measured at 200 mM NaCl (N = 1019). The inset represents the difference between the measured and the fitted 2D density profile, i.e. the residuals of the fit. The occupancies determined from the fits are shown in Fig. 6c.

**a**

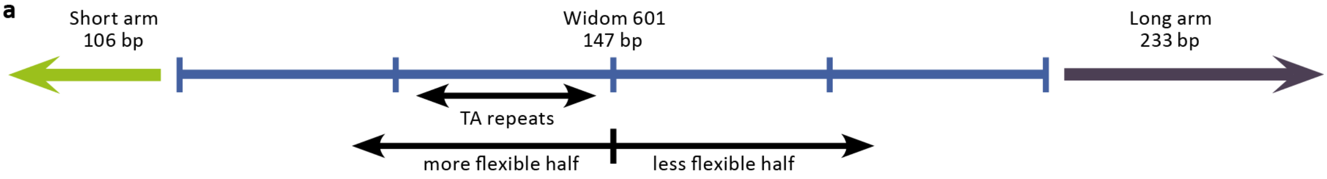


CAATCAAACTGGCTCGTCGCGAATTGGAGCTCCACCGCGGTGGCGGCCGCTCGATCTAGTACTAGTGGCATGTCAGCTGCAGGAATTCGAGCTCAACGTG
CAATCCCTGGAGAATCCCGGTGCCGAGGCCGCTCAATTGGTCGTAGACAGCTCTAGCACCGCTTAAACGCACGTACGCGCTGTCCCCCGCGTTTTAACCG
CCAAGGGGATTACTCCCTAGTCTCCAGGCACGTGTCAGATATATACATCCTGTCGACGACACGGGTGATCGACTAGTTCTAGAGCGATCTAGTATCGATC
ACTCTTTTGTTCCCTTTAGTGAGGGTTAATTTCGAGCTTGCGATCTAGTCGATACTACGCGTAATCATGGTCATAGCTGTTTCCTGTGTGAAATGATCTA
CTTGTTATCCGCTCACAATTCCACACAACATACGAGCCGGAAAAGAAATAAAGATCTAGACTACTGCATAAAGTGTAAAGCCTGGG

**Supplementary Fig 4 | Nonpalindromic nature of the W601 positioning sequence**.

**a**, The 486 bp DNA construct contains the Widom 601 nucleosome positioning sequence (147 bp) flanked by a short arm (106 bp) and a long arm (233 bp). Four TA repeats with a 10 bp periodicity on the short arm half of the W601 sequence induce the higher flexibility of the half of the W601 sequence on the short arm side[17].
